# Network-based Computational Drug Combination Prediction

**DOI:** 10.1101/049015

**Authors:** Fuhai Li, Huang Lei, Jianting Sheng, Stephen Wong

## Abstract

Cancers are complex diseases that are regulated by multiple signaling pathways. Patients often acquire resistance to single drug treatment. Use of drug combinations that target multiple parallel pathways is a promising strategy to reduce the drug resistance. Pharmacogenomics big data are being generated to uncover complex signaling mechanisms of cancers and correlate cancer-specific signaling with diverse drug responses. Thus, converting pharmacogenomics big data into knowledge can help the discovery of synergistic drug combination. However, it is challenging and remains an open problem due to the enormous number of combination possibilities and noise of genomics data.

## I. Introduction to the type of problem in cancer

Cancers are complex diseases regulated by interactions of multiple signaling pathways. Though a number of anti-cancer drugs are currently in use, they not are curative and many are limited by acquired drug resistance. Alternative signaling pathways can maintain tumor development. Use of drug combinations that target multiple parallel pathways [1] is a promising strategy to reduce the drug resistance effect, and improve cancer treatment outcomes. However, it is infeasible to discover drug combinations experimentally due to the enormous number of combination possibilities.

Recent years have seen the explosive generation of large volumes of pharmacogenomics data with the primary goals to uncover the complex signaling mechanisms of cancers and correlate cancer-specific signaling with diverse responses to thousands of drugs. For example, the cancer genome atlas (TCGA) project profiles the genomics of over 10,000 patient samples across over 20 types of cancer [2, 3]. Integrative analyses often indicate multiple subtypes of cancers with complex genomics characteristics [4, 5]. The Cancer Cell Line Encyclopedia (CCLE) [6], Genomics of Drug Sensitivity in Cancer (GDSC) [7, 8]. The Connectivity Map (CMAP) [9] generated gene expression signature data of different cell lines under various perturbations by 1,039 small molecules. These data have since been scaled over 1,000 times (including 5585 drugs and bioactives, and 20,000+ genetic reagents (targets)) and are publicly available at (http://www.lincscloud.org/) (LINCS: Library of Integrated Network-based Cellular Signatures).

Though rich information is embedded in the pharmacogenomics big data, it remains an open problem to convert that big data to knowledge for discovery drug combination. Some of the challenges are as follows. First, there are millions of possible pair-wise drug combinations of even 1000 drugs. Secondly, the genomics profile data are noisy for both cancer patients, and compounds and genetic reagents from CMAP; and 2) the selection of synergistic drug combinations from millions of possible combinations is difficult. Therefore, novel data-driven approaches are needed urgently to integrate genomics profiles of cancers and thousands of drugs and molecules, and genetic reagents to predict the synergistic drug combinations.

Network-based drug discovery is touted as the next drug discovery paradigm [10]. Diseases are often regulated by complex signaling networks, and multi-drugs and multi-targets are often associated to form a big drug-drug, and drug-target network. On the other hand, there are rich computational resources to support the network analysis and visualization. In this study, we introduce a network-based computational approach for synergistic drug combination prediction [11].

## II. Illustrative Results of application of Methods

The proposed drug combination prediction approach was evaluated on lung adenocarcinoma. The genomics profiles of lung adenocarcinoma patients were collected from GEO (Gene Expression Omnibus) (GDS3257) to reconstruct the lung adenocarcinoma-specific signaling network. The genomics profiles of 1309 drugs and compounds were obtained from CMAP to build the drug network modules. Three drugs, Gefitinib, Paclitaxel and LY-294002, were used as the anchor drugs, and the optimal drugs combining with them are predicted (ranked). Through literature review as the validation, among the top 50 predicted drug pairs, 9, 8, and 3 drug pairs have been reported to be effective respectively [11]. This success rate is significantly higher compared with other drug combination methods and random selection [11]. Figure 1 shows the distribution of Gefitinib and Paclitaxel targets on the reconstructed lung adenocarcinoma disease signaling network, which indicates the potential mechanism of action of the drug combination.

**Figure 1.**
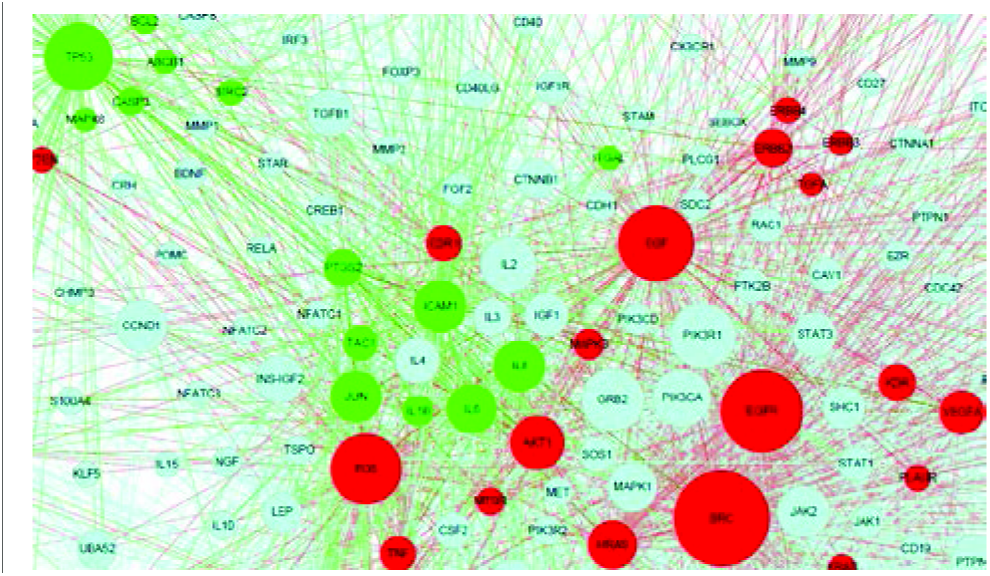
Target distribution of (red) Gefitinib and (green) Paclitaxel on the reconstructed lung adenocarcinoma dsease signaling network.

## III. Quick Guide to the Methods

### A. Major assumptions of the model

The proposed approach was designed based on the following major assumptions. First, disease signaling networks have alternative signaling modules. Secondly, drugs can inhibit multiple targets on the disease signaling network. Thirdly, effective drug combinations can inhibit alternative signaling modules simultaneously.

### B. Methodology Overview

Figure 2 shows the methodology overview. In general, the drug-drug similarity (distance) is estimated based on the genomics profiles of drugs from the CMAP database [12]. Other information can also be integrated to refine the drug-drug similarity, e.g., the drug-target interaction information in STITCH [13]. Then drugs are connected into a drug-drug network given a similarity threshold, which is partitioned into drug network modules using the network analysis [12] or clustering approaches [11]. Different drug network modules might share the common mechanism of action, and their targets can indicate the inhibited signaling modules of diseases. The drug target information can be obtained from the following database, e.g., DrugBank [14], STITCH [13], NPC browser [15]. The disease signaling network can be reconstructed [16, 17] by integrating disease genomics data and protein interactome data, e.g., BioGRID [18], Biocarta [19], Reactome [20], HPRD [21], and KEGG [22]. Drugs (from different drug network modules) are combined to disrupt disease signaling network modules using the network centrality metrics [23, 24].

**Figure 2:**
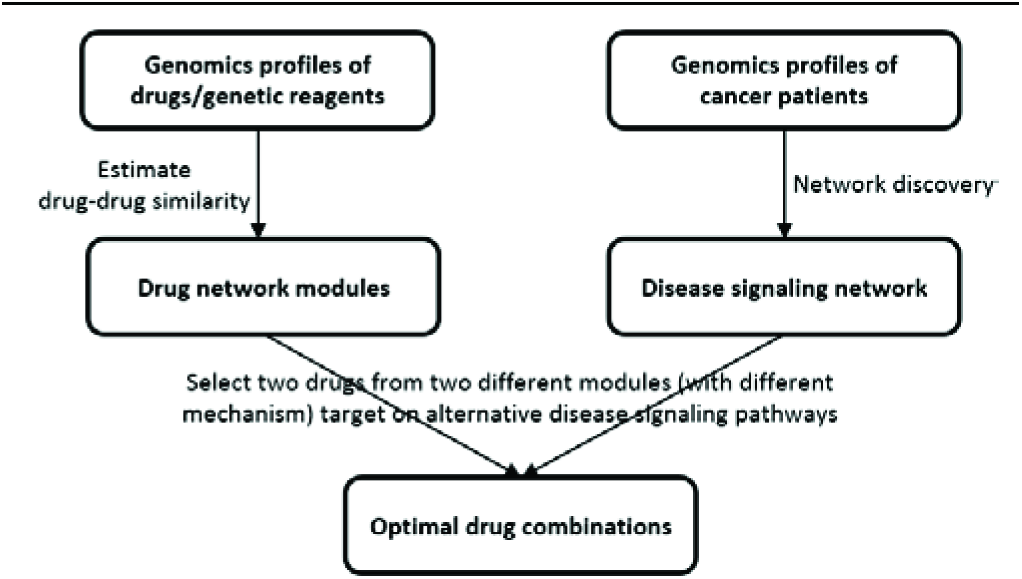
Methodology Overview

### C. Network analysis and visualization tools

There are rich computational resources of network-based approaches that can be used for biomarker and drug discovery. Some widely used network analysis tools are, e.g., igraph in R (http://igraph.org/r/), NetworkX in Python (https://networkx.github.io/) for network structure analysis, and, e.g., Cytoscape (http://www.cytoscape.org/), and D3.js (http://d3js.org/) for network visualization.

## Acknowledgment

We would like to thank the bioinformatics group of the department of Systems Medicine and Bioengineering for their helpful discussions.

